# BMP signalling attenuates intercellular adhesion to drive mesenchyme migration during fin fold morphogenesis

**DOI:** 10.1101/2024.08.18.608445

**Authors:** Bitan Saha, Harsha Mahabaleshwar, Charmaine Ho Min, Leslie Boon Haw Leong, Levene Wenqian Chua, Samuel Kwok, Tom J. Carney

## Abstract

The development of the vertebrate limb bud is a multifactorial process involving the coordinated activity of diverse signalling pathways and cell types. BMP signalling has been consistently shown to play an integral role during the patterning of mouse and chick limbs. However, the molecular mechanisms and cellular processes underlying its contribution to limb development are poorly understood. Using zebrafish embryos, we demonstrate a novel role of BMPs in driving median fin fold morphogenesis, an evolutionary related structure. Our findings indicated that a graded BMP signal is established in the emerging fin fold along the proximo-distal axis, with the fin mesenchymal cells being identified as the primary target of the signalling cue. Pharmacological and genetic ablation of BMP signalling in the fin fold inhibited the normal migration of the mesenchymal cells into the fin fold. The observed inhibition of migration was accompanied by the elongation in mesenchymal cell shape, which hinted at the possibility of altered physical forces acting on the cell population. We observed that loss of BMP signalling led to the cellular re-distribution and altered membrane dynamics of N-cadherin in the fin mesenchymal cells, which manifested as an increase in homotypic cell adhesion. This change in cell-adhesion serves as the mechanistic basis for the observed defect in migration. Furthermore, inability to detach due to loss of BMP signalling occurs independently of sphingosine-1-phosphate signalling that generates a gradient of mesenchyme adhesion to the underlying matrix. Overall, this study highlights the cellular and molecular mechanism underlying the role of BMPs in coordinating early fin bud morphogenesis.

## Introduction

Animal development is composed of a sequence of intricate biochemical and cellular events that are temporally orchestrated by a central group of molecules that provide spatial information to the embryonic cells to coordinate tissue patterning and influence fate decisions [1, 2]. These ‘morphogens’ include signalling factors such as the Bone Morphogenic Proteins (BMPs) that are indispensable for embryogenesis, with deleterious mutations in these genes and their regulatory partners often manifesting as a plethora of lethal morphological defects and severe functional disorders [3–5]. Although more frequently associated with the early stages of development, morphogens have also been recognised to play pivotal roles in later stages, encompassing biological processes such as organogenesis, tissue growth and homeostasis.

BMPs belong to the TGF-β family of growth factors with implications in a myriad of cellular processes including but not limited to specification, proliferation, differentiation and apoptosis [6–10]. Similar to other members of the protein superfamily, signalling by BMPs are initiated by the binding of mature dimeric ligands to its cognate cell surface receptor, and the formation of a heterotetrameric complex comprising both Type I and Type II BMP receptors [11]. The activated receptor complex phosphorylates substrate proteins called R-Smads (e.g. Smad1/5/9) which forms complexes with Co-Smads and translocate to the nucleus where they regulate the transcription of downstream target genes [12]. In addition to the Smad-mediated canonical pathway, BMPs have been shown to signal through several Smad-independent non-canonical pathways [13]. These pathways utilize signal transducers such as ERK, p38 MAPK, PI3K/Akt, Rho-GTPases and LIMK among others [14–23]. The choice between the canonical and non-canonical pathways is determined by several factors including nature of Type I receptor involved and the oligomerization state of the BMP receptor complex [24]. However, owing to the convoluted and oftentimes overlapping nature of the two pathways, it is often difficult to ascertain which pathway is responsible for a particular biological action.

The morphogenetic role of BMPs are well documented in several model organisms where they have been shown to govern embryonic patterning along the dorsoventral axis and direct the specification of mesodermal cell fates [7, 25, 26]. In addition to early embryogenesis, BMP signalling has been shown to be involved in the development of several major organs and tissue systems such as muscles, bones, heart, vasculature, lungs, kidney, eyes, limb bud and the gastrointestinal tract [27–33]. Of particular interest is the multifaceted role of BMPs in vertebrate limb morphogenesis which has been extensively studied using murine models [34]. Studies have demonstrated the presence of BMPs in the developing limb ectoderm where it regulates limb outgrowth and controls digit formation and identity [35, 36]. Conditional knockout of BMP receptors in the limb ectoderm displayed a range of patterning defects including amelia and polysyndactyly [37]. Similar observations were reported upon abrogation of BMP signalling in the murine limb mesoderm [38, 39]. The fundamental importance of this pathway in limb development is reinforced by concurrent studies in other model systems, where BMPs and their orthologues have been shown to be crucial for the proper growth and morphogenesis of the Drosophila wing disc and zebrafish pectoral fin [40, 41]. However, the cellular and molecular mechanisms behind the role of BMP signalling in limb development are poorly understood.

Here, we explore a novel function of BMP signalling in the morphogenesis of the zebrafish larval fin fold, a precursor to the adult median fins. During the early stages of fin fold development, we observed the presence of a BMP ligands in the fin ectoderm which temporally coincided with the expression of BMP receptors in the fin mesoderm. This signalling relay between ectoderm and mesoderm helps to establish a BMP activity gradient in the cells of the fin mesenchyme along the proximo-distal axis. Disruption of this activity gradient, either by pharmacological or genetic methods, inhibited the usual migration of the fin mesenchyme cells across the fin fold and impeded proper fin growth. Morphometric analysis of mesenchyme cells using transgenic fish demonstrated significant changes in morphology, in parameters such as circularity and cellular arborisation. An investigation into the expression and dynamics of cell adhesion molecules revealed changes in the cellular distribution and membrane mobility of N-cadherin in fin mesenchyme cells that had been subjected to BMP inhibition, hinting at a possible mechanism by which BMP signalling orchestrates cell migration via regulation of N-cadherin-mediated intercellular adhesion.

## Results

### A BMP signalling gradient is established during median fin fold development in zebrafish

In mice, BMP ligands and receptors have been shown to be expressed in both the ectoderm and mesoderm of the developing limb bud [42], and limb malformations were observed when BMP signalling was abrogated in either the limb ectoderm or mesenchyme [36, 39, 43], suggesting a tight interplay in BMP signalling between mesoderm and ectoderm. With this in mind, we sought to explore the role of BMP signalling in zebrafish limb development using the median fin fold (MFF) as the model tissue system. In situ hybridization (ISH) revealed the expression of major BMP ligands such as *bmp2b*, *bmp4* and *bmp7b* in the apical ectodermal ridge (AER) of the MFF at 24 hpf and 36 hpf (Figure 1A-C, A’-C’), which had mostly disappeared by 48 hpf (Figure 1A’’-C’’). BMP Type 1 receptors such as *bmpr1ab* and *bmpr1ba* were also expressed at similar developmental stages but were restricted to the fin mesoderm (Figure 1 D-D”, E-E”). Surprisingly, even though expression of BMP ligands and receptors could not be detected in the MFF beyond 36 hpf, we observed strong expression of *id1*, a downstream target of BMP signalling [44], at all tested stages throughout the fin mesenchyme (Figure 1 F-F”).

**Figure 1.**
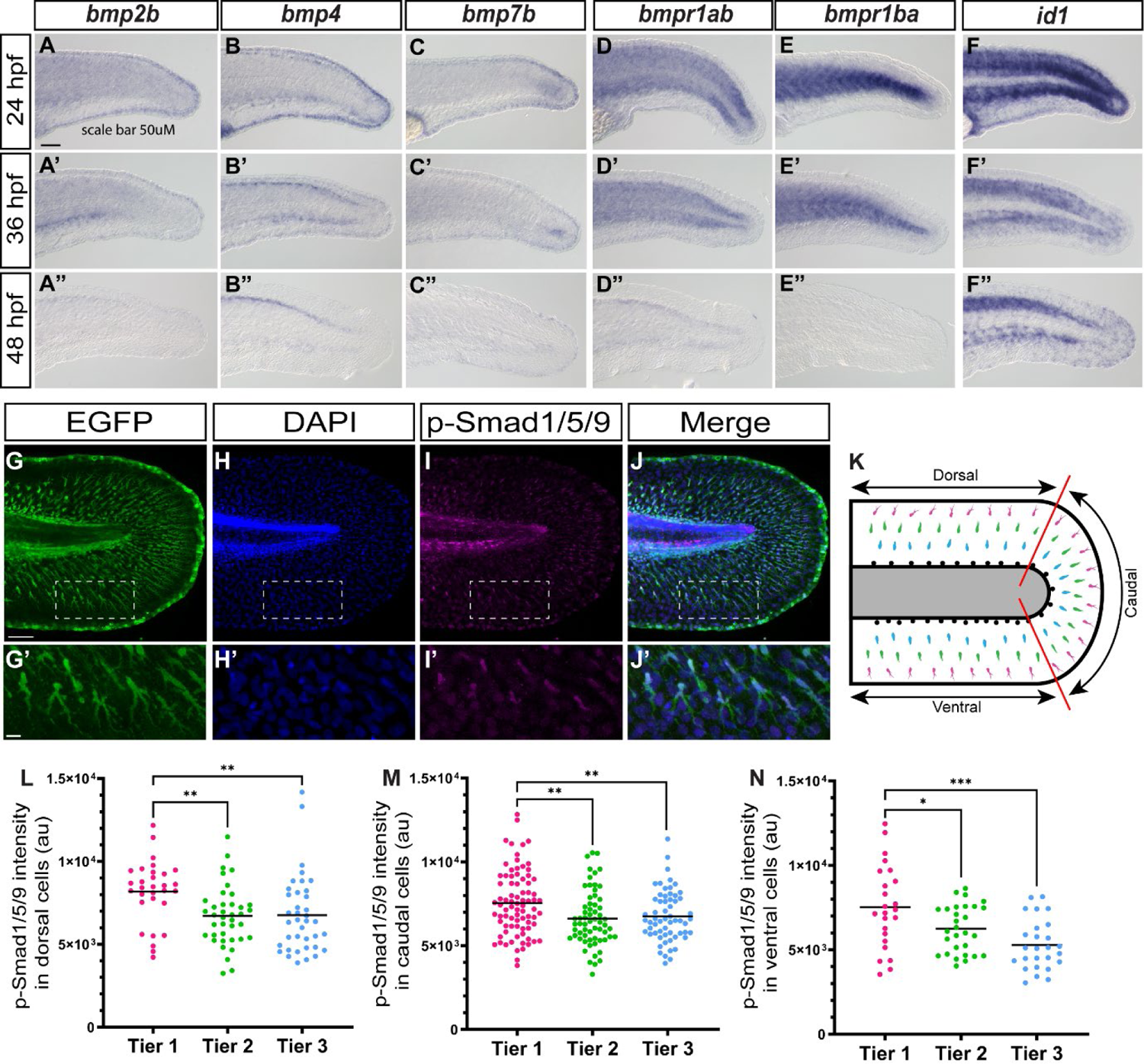
A BMP activity gradient is established in the developing fin fold. In situ hybridization revealed the expression of BMP ligands, receptors and downstream targets in the developing fin at 24 hpf (A-F), 36 hpf (A’-F’) and to some degree at 48 hpf (A”-F”). Scale bar represents 50 µM. IHC with pSmad-1/5/9 antibody demonstrated BMP activity in the fin fold (G-J) with particularly high expression in the cells of the fin mesenchyme (G’-J’). For quantification purposes the fin fold was divided into sections and the mesenchyme cells were categorized into tiers (K). Quantification of nuclear pSmad-1/5/9 intensity demonstrated the presence of a BMP activity gradient along the proximo-distal axis of the fin fold (L-N).

Having observed expression of *id1* transcripts in the fin mesenchyme, we sought to confirm whether the cells of the fin mesenchyme might be responsive to the BMP signalling cue by direct *in vivo* imaging of the cell population. To do so, we utilized a transgenic enhancer trap line ET37, in which fin mesenchymal cells express GFP [45] in conjunction with a BMP reporter line BRE:dmKO2, that expresses destabilized mKO2 under the regulation of a BMP response element (BRE) and has been shown to provide a dynamic readout of Smad-mediated BMP signalling [46]. We found that between 24 hpf and 48 hpf, reporter activity could be detected in the GFP^+^ cells of the fin mesenchyme that had delaminated from the mesoderm and emigrated outwards to colonize the larval fin (Supplemental Figure 1A, Supplemental Movie 1) [47, 48]. We also observed that at 48 hpf, the fin mesenchyme cells that were closer to the AER appeared to have stronger BMP activity compared to the ones farther away, indicating the possibility of a BMP activity gradient along the proximo-distal axis.

To confirm the existence of an active BMP signalling gradient we performed immunohistochemistry (IHC) for phosphorylated Smad1/5/9 in ET37 embryos at different stages of fin fold formation. We observed robust pSmad-1/5/9 signal in the median fin fold at 24 hpf, 36 hpf and 48 hpf (Figure 1G-J, Supplemental Figure 1B). Similar to our observations with the BMP reporter transgenic line, the pSmad-1/5/9 staining was enriched in the nucleus of the migrating fin mesenchymal cells (Figure 1G’-J’). To simplify the quantification of the signalling gradient across the cells of the fin fold, we divided the fin fold at 48 hpf into three distinct sections based on anatomical position – dorsal, ventral and posterior, and within each section we sub-divided the cell population into three concentric tiers based on their position along the proximo-distal axis (Figure 1K). We found that across all sections, Tier 1 cells (distally positioned) had significantly higher levels of nuclear pSmad1/5/9 when compared to Tier 2 cells (intermediate position) and Tier 3 cells (proximally positioned), providing evidence for the existence of a BMP activity gradient along the P/D axis (Figure 1L-N).

### Inhibition of BMP signalling in the fin fold affects fin mesenchyme migration

In order to investigate the physiological role of BMP signalling in the fin fold, we used small-molecule antagonists of BMP Type I receptors such as K02288 [49] and LDN193189 [50] to disrupt the signalling pathway in the MFF. Embryos treated with either K02288 or LDN193189 from 28 hpf to 48 hpf exhibited reduced migration of the fin mesenchyme cells into the fin fold compared to embryos treated with vehicle control (Figure 2 A-A”, B-B”, Supplemental Movie 2. A robust phenotype was observed upon treatment with 25 µM of either inhibitor and further increasing the concentration did not have any significant effect on phenotype severity (Figure 2 C, D). Though both antagonists had equivalent effects on fin mesenchyme migration, embryos treated with higher concentrations of LDN193189 showed reduced survival rates over extended treatment periods (data not shown).

**Figure 2.**
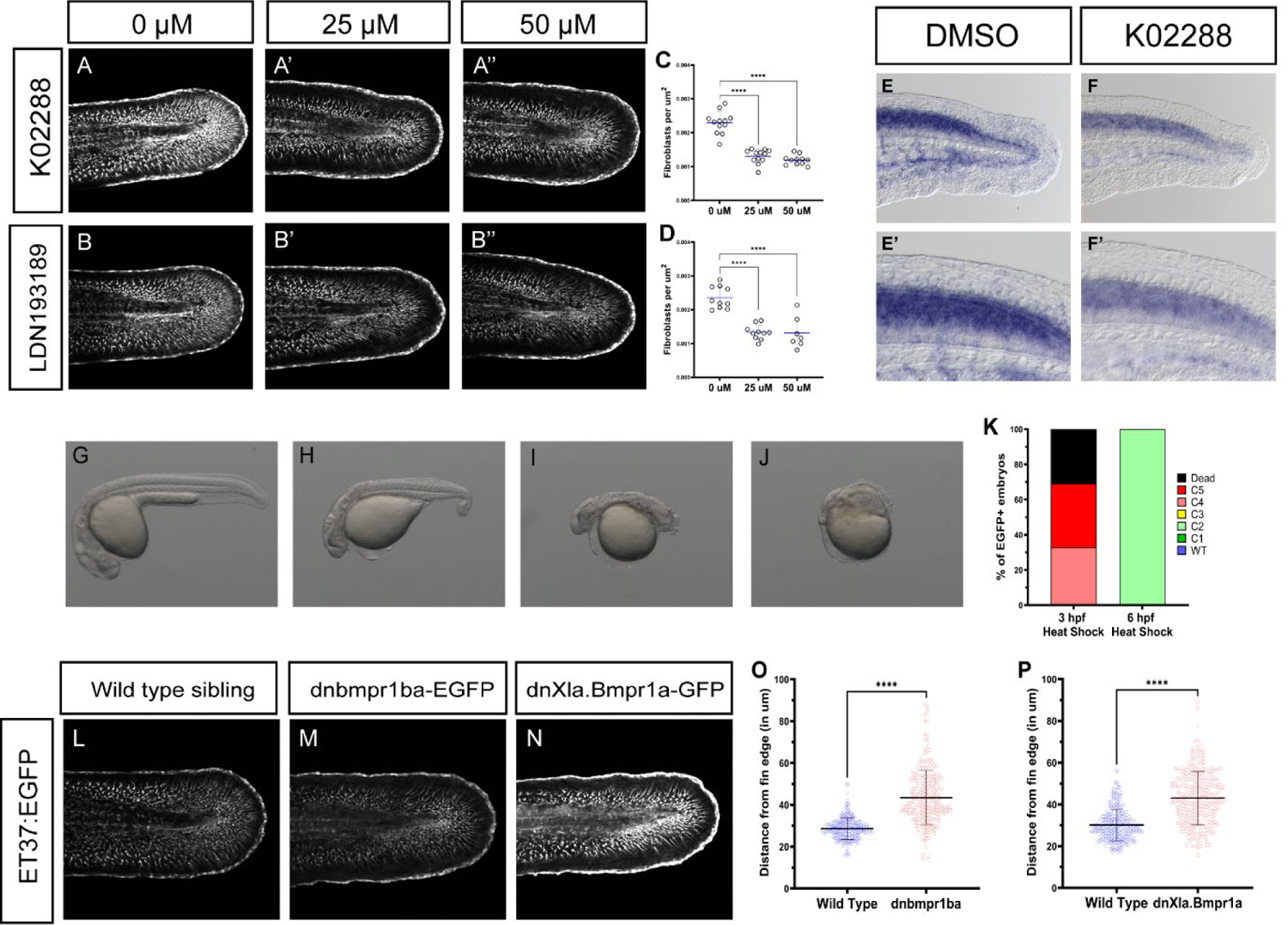
Inhibition of BMP signalling affects the migration of the fin mesenchyme. Treatment of ET37:EGFP embryos with increasing concentration of small-molecule BMP antagonists such as K02288 (A-A”) or LDN193189 (B-B”) results in the retarded migration of fin mesenchyme cells. A strong phenotype is observed at 25 µM with further increase not resulting in any significant changes (C, D). In situ hybridization shows decreased levels of *id1* transcript in the fins treated with K02288 (E, F) with the strongest reduction observed in the cells of fin mesenchyme that show migration defects (E’, F’) suggesting a causal relationship between BMP activity and cell migration. Transgenic Tg(*hsp70l:dnbmpr1ba-EGFP,crybb:ECFP*) embryos demonstrated a range of dorsalization phenotypes (H-J) compared to wild-type siblings (G) when heat-shocked during gastrulation. More severe phenotypes were observed when heat-shock was performed at 3 hpf as compared to 6 hpf (K). Similar to the treatment with K02288, fin mesenchyme migration was affected in both transgenic lines over-expressing dominant-negative BMP receptors (L-N). Quantification of cell migration showed similarly significant effects in both transgenic lines (O, P).

Next, embryos treated with either K02288 or vehicle control were subjected to whole mount ISH with an anti-sense probe against *id1*. The results showed a global reduction in the level of *id1* transcript across the median fin fold of embryos treated with K02288, concurrent with the expected reduction of BMP activity (Figure 2 E, F). An evident reduction in *id1* transcript levels were observed in the fin mesenchymal cells that had failed to detach from the mesoderm and had not migrated out into the fin fold upon BMP inhibition (Figure 2 E’, F’). These observations indicate the possibility that migration of the fin mesenchyme may be dependent on the BMP activity levels in the migrating cell population. The correlation between loss of BMP activity in the fin mesenchyme cells and the loss of migratory behavior was further validated using IHC (Supplemental Figure 2A, B).

In addition to the pharmacological inhibition of BMP signalling we validated our observations by transgenic overexpression of a dominant-negative BMP receptor. We generated the Tg(*hsp70l:dnbmpr1ba-EGFP,crybb:ECFP*) line that could conditionally repress BMP signalling by transient overexpression of a truncated zebrafish Type I BMP Receptor (BMPR1ba) under the regulation of the heat-inducible *hsp70* promoter. Similar to a previously described dominant-negative transgenic line Tg(*hsp70l:dnXla.Bmpr1a-GFP*) derived from *X.laevis* BMP receptor [51], our dnbmpr1ba-EGFP embryos exhibited varying degrees of dorsalization when heat-shocked during early development (Figure 3 G-J). The degree of dorsalization correlated with the onset of transgene expression, with the most severe phenotypes displayed by embryos where transgene expression was initiated prior to the onset of gastrulation (Figure 3K). When the dnbmpr1ba-EGFP transgene was overexpressed in ET37 embryos, migration defects were observed in the cells of the fin mesenchyme identical to those seen upon treatment with BMP antagonists (Figure 2L, M). Identical results were also obtained in experiments with the Tg(*hsp70l:dnXla.Bmpr1a-GFP*) transgenic embryos (Figure 2N). The changes in cell migration were accompanied by a significant reduction in BMP reporter activity in both transgenic lines (Supplemental Figure 2C, D), further suggesting a causal relationship between BMP signalling and fin mesenchyme migration.

**Figure 3.**
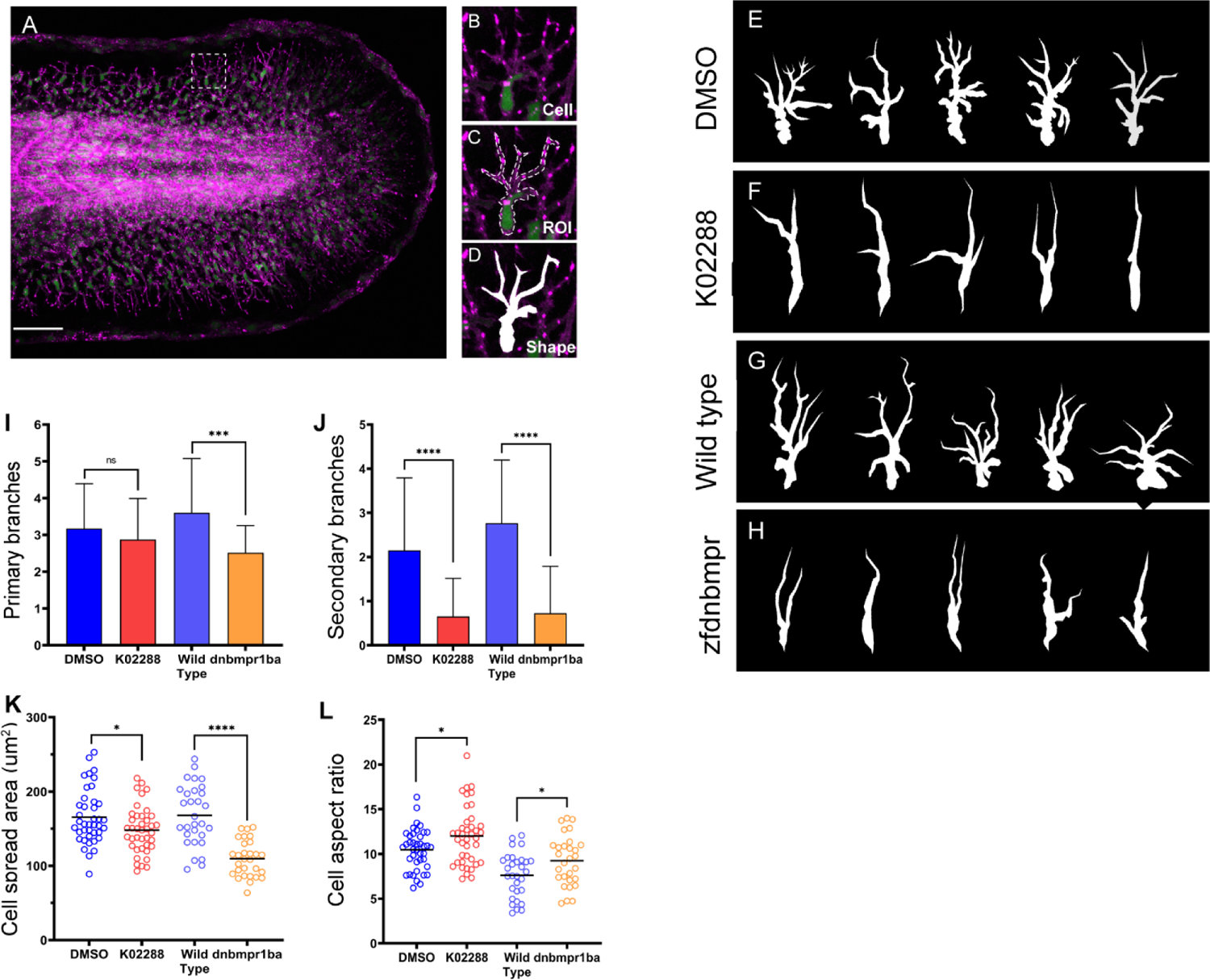
Inhibition of BMP signalling affects cellular morphology of fin mesenchyme cells. The transgenic line Tg(*tbx16l:nlsGFP-P2A-mCherryCAAX*) co-expressed nuclear GFP and farnesylated mCherry in cells of the fin mesenchyme (A, B) allowing us to trace the outer boundaries of the cell (C) and extract the cell shape (D). Comparing the shape traces of cells from embryos treated with vehicle control (E) and K02288 (F) as well as wild-type embryos (G) and siblings overexpressing the dnbmpr1ba-EGFP transgene (H), we were able to demonstrate observable differences in cellular morphology under conditions of BMP signalling inhibition. The most notable difference was the significant reduction in the number of primary (I) and secondary (J) cellular branches. We also observed a decrease in cell spread area (K) and an increase in cell aspect ratio (L) upon BMP inhibition.

The signalling cascade downstream of activated BMP receptors may be transduced through different effector molecules depending on the activation of the canonical or non-canonical pathways [12, 13]. To examine the individual contributions of these distinct molecular networks, we generated a transgenic line Tg(*hsp70l:smurf1-P2A-mCherryCAAX, crybb:ECFP*) that enables selective abrogation of the Smad-mediated canonical pathway by transiently over-expressing Smurf1, an E3 ubiquitin ligase that targets BMP-specific R-Smads for degradation via ubiquitination [52, 53]. Transgene activation induced dorsalization of embryos during gastrulation and led to reduced expression of *id1* transcripts in the fin fold, demonstrating the BMP-inhibitory activity of Smurf1 (Supplementary Fig 3A-C). However, Smurf1 over-expression did not have any significant effect on fin mesenchyme migration (Supplementary Figure 3D-E), indicating that the canonical pathway may not have major contributions to the pro-migratory activity of BMP in the developing median fin fold. An alternate explanation for our observations might be that owing to the transient nature of the over-expression, Smurf1 protein levels were not maintained at adequate levels throughout the entirety of the migratory period resulting in an insufficient repression of the pathway.

In cell-specific contexts, BMPs are capable of exerting biological action through secondary signalling effectors such as MAPKs and other associated kinases. We assessed the possible contributions of two such non-canonical effectors of BMP signalling, namely ERK and p38, in driving fin mesenchyme migration. However, we did not observe any significant changes in ERK or p38 activity in mesenchymal cells whose migration had been impeded by BMP inhibition (Supplementary Figure 4 A, B). Taken together, our findings provide a limited yet valuable insight into the signalling landscape of fin mesenchyme during the early stages of fin development.

### The migration defect is accompanied by changes in cellular morphology

Initial observations of fin mesenchyme cells under conditions of BMP signalling inhibition suggested changes in cellular appearance. To validate this hypothesis, a transgenic line Tg(*tbx16l:nlsGFP-P2A-mCherryCAAX*) was generated that would allow for the distinct visualization of cellular boundary and nuclear position by co-labeling of fin mesenchymal cells with nuclear targeted GFP and plasma-membrane localized mCherry (Figure 3A, B). Using the labelling with mCherry, the outer boundaries of these cells were manually traced (Figure 3C) and the traced region of interest was filled to extract the cell shape (Figure 3D). At 48 hpf, fin mesenchymal cells in DMSO treated control embryos were seen to be highly arborized with several primary projections extending apically from the cell body and each primary projection giving rise to one or more secondary projections. In contrast, fin mesenchymal cells in K02288 treated embryos displayed strongly reduced arborization with fewer noticeable projections per cell (Figure 3E, F). Similar trends were also observed in fin mesenchymal cells of embryos with dominant-negative BMPR1ba overexpression when compared to their wild type siblings (Figure 3G, H). We also noted a subtle morphological change at the rear end of the migrating mesenchymal cells wherein the trailing edge under control conditions appeared more rounded compared to the pointed trailing edge observed in BMP inhibited embryos (Figure 3E-G).

We followed up our observations by conducting a morphometric analysis of the fin mesenchymal cells to gain quantitative insight into these cellular changes. The most prominent finding was a significant decrease in the degree of arborization, i.e. the number of primary and secondary cellular projections, in samples that were subjected to either pharmacological or genetic repression of BMP signalling as compared to their un-repressed counterparts (Figure 3I, J). Cells under BMP repression also exhibited a significantly reduced cell spread area (Figure 3K) as well as an increased cell aspect ratio (Figure 3L) suggesting an elongated, spindle-like appearance. These results hinted at a possibility that in addition to a reduction of migratory behavior, the inhibition of BMP signalling may also lead to the generation of localized physical forces that act upon the mesenchymal cells to distort their morphology.

### BMP-dependent mesenchyme migration may be mediated by changes in cell adhesion dynamics

In order to understand the molecular mechanisms underlying mesenchymal cell migration, we studied fin fold growth in real-time at high temporal resolutions to identify subtle changes in cellular behavior in the absence of BMP signalling. Time-lapse imaging of ET37 embryos under physiological conditions revealed that starting at around 28 hpf, the fin mesenchymal cells progressively separated from the fin mesoderm and migrated towards the AER. Along the path, these cells often came into contact with neighboring migrating cells, however these interactions were always transitory, lasting only a few minutes before the cells diverged (Figure 4A, Supplemental movie 3). In sharp contrast, the majority of fin mesenchyme cells under K02288 treatment failed to disengage from the mesoderm and initiate migration. Cells that had already separated from the mesoderm prior to the addition of inhibitor continued to migrate until they made contact with stationary neighboring cells, at which point their movements practically ceased. Under these conditions, contact period between cells were prolonged and the cells continued to extend while remaining attached to their neighbors, giving rise to chains of elongated cells (Figure 4A, Supplemental movie 3). Trajectory projection of selected cells also revealed that under K02288 treatment mesenchymal cells had travelled significantly less distance from their origin and had correspondingly reduced average speed (Figure 4B, C). However, unlike Slit-Robo signalling whose disruption affects the polarity of the fin mesenchymal cells, abrogation of BMP signalling did not appear to have any effect on cell directionality (Supp rose plot).

**Figure 4.**
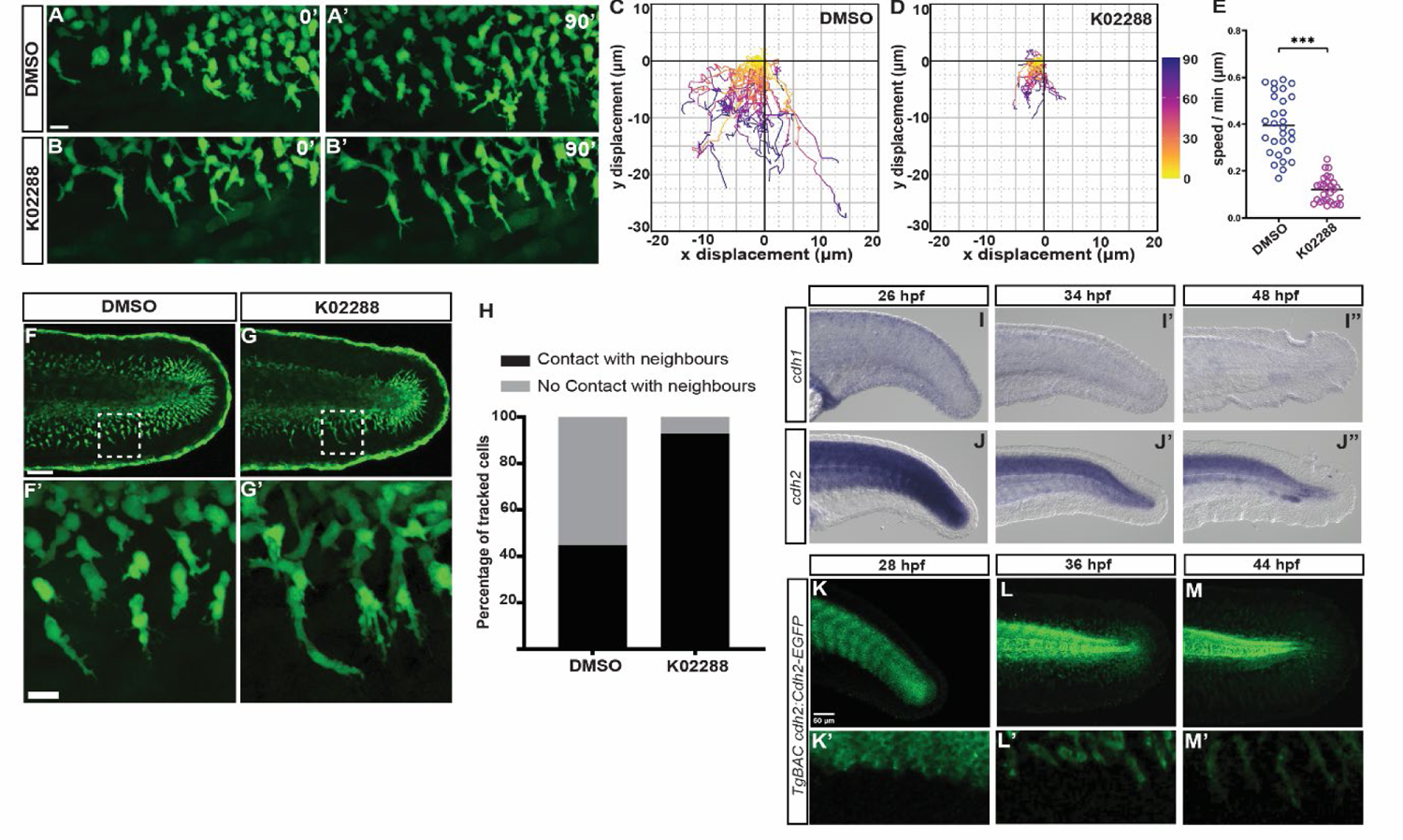
Attenuation of fin mesenchyme migration may be attributed to changes in cell adhesion. Snapshots from the confocal time lapse of migrating fin mesenchyme highlight the difference in migratory behavior between DMSO-treated (A, A’) vs. K02288-treated (B, B’) embryos. Plotting the trajectory of selected cells allows us to visualize the difference in migratory path (C, D) as well as the migration speed (E) under these conditions. Subsequent analysis of high–resolution images of the developing fin fold in control (F, inset F’) and BMP-inhibited conditions (G, inset G’) demonstrate the propensity of mesenchymal cells to form interconnected chains in the absence of BMP signaling. Quantification of cell-cell contact events at the end of 90’ interval (tracked mesenchymal cells analyzed in A-E) demonstrate a significant increase of homotypic attachment in the absence of BMP signaling (H). In situ hybridization reveals the expression of *cdh2* (J-J”) but not *cdh1* (I-I”) in the fin mesenchyme cells between 26 hpf and 48 hpf. Imaging of TgBAC(*cdh2:cdh2-EGFP*) embryonic fins between 24 hpf and 48 hpf show the dynamic expression of Cdh2-EGFP in the fin mesenchyme cells (K-M). As the cells migrate further into the fin fold, the intensity of Cdh2-EGFP in the cells appear to decrease with the proximal cells retaining the highest levels of Cdh2-EGFP and the distal cells the lowest (K’-M’).

To gain a quantitative measure of the degree of cell attachment, we further examined the selected cells from our time-lapse images to compare the proportion of cells having physical contact with their neighbors under different conditions of BMP signalling. We found that in contrast to control conditions which had a majority (>50%) population of free-floating cells at the end of the tracking period, almost all the mesenchymal cells in K02288 treated embryos exhibited some degree of contact with their neighbors (Figure 4D). Furthermore, we observed that upon treatment of embryos with K02288 and subsequent washout, the increase in intercellular contact was reversed as the population of free-floating migratory cells was largely restored (Supplemental movie 4). This led us to hypothesize that the observed defect in migration may perhaps be a derived effect of the fin mesenchymal cells undergoing changes in homotypic cell adhesion in the absence of BMP signalling.

BMPs are known to affect cellular movement in several cell types via regulation of classical homotypic cellular adhesion molecules such as E-cadherin and N-cadherin [54, 55]. ISH revealed the expression of both E-cadherin (*cdh1*) and N-cadherin (*cdh2*) transcripts in the fin fold. However, where *cdh2* transcripts could be detected in the cells of the fin mesenchyme, *cdh1* was only expressed in the epidermal cells of the fin fold (Figure 4E), suggesting a higher probability of N-cadherin involvement. Using the BAC transgenic line TgBAC(*cdh2:cdh2-EGFP,crybb1:ECFP*) expressing a Cdh2-EGFP fusion driven by the endogenous *cdh2* promoter [56], we studied the cellular distribution of N-cadherin in the fin mesenchyme cells during their migration into the fin fold. Live imaging across multiple developmental stages revealed higher levels of N-cadherin expression in the proximal mesenchymal cells with expression progressively decreasing as the cells migrated further into the distal fin fold (Figure 4F, Supplemental Movie 5). This indicated that reduction of cellular N-cadherin levels might be a requirement for the mesenchymal cells to successfully colonize the fin fold.

The downregulation of N-cadherin in actively migrating cells was suggestive of a possible correlation between BMP signalling and N-cadherin expression. Time lapse microscopy of TgBAC(*cdh2:cdh2-EGFP,crybb1:ECFP*)embryos under different conditions of BMP signalling demonstrated major changes in cellular distribution of N-cadherin in the fin mesenchyme. Under physiological conditions, Cdh2-EGFP was mostly uniformly distributed on the plasma membrane of the fin mesenchymal cells, with the expression levels gradually seen decreasing with time as the cells detached from the mesoderm and moved farther into the fin fold (Figure 5A, Supplemental Movie 6). In the absence of BMP signalling, Cdh2-EGFP in the mesenchymal cells appeared to cluster together into punctate structures on the plasma membrane (Figure 5B,Supplemental Movie 6). Though these puncta could be observed occasionally under control conditions, the frequency of puncta per cell was drastically higher under BMP inhibition (Figure 5C). A similar redistribution of N-cadherin was also observed upon BMP inhibition with LDN193189 (Fig 5D, E). BMP-inhibited embryos also exhibited no changes in fin mesenchymal E-cadherin levels, further cementing the noninvolvement of E-cadherin in modulating mesenchymal cell adhesion (Supplemental Figure 5).

**Figure 5.**
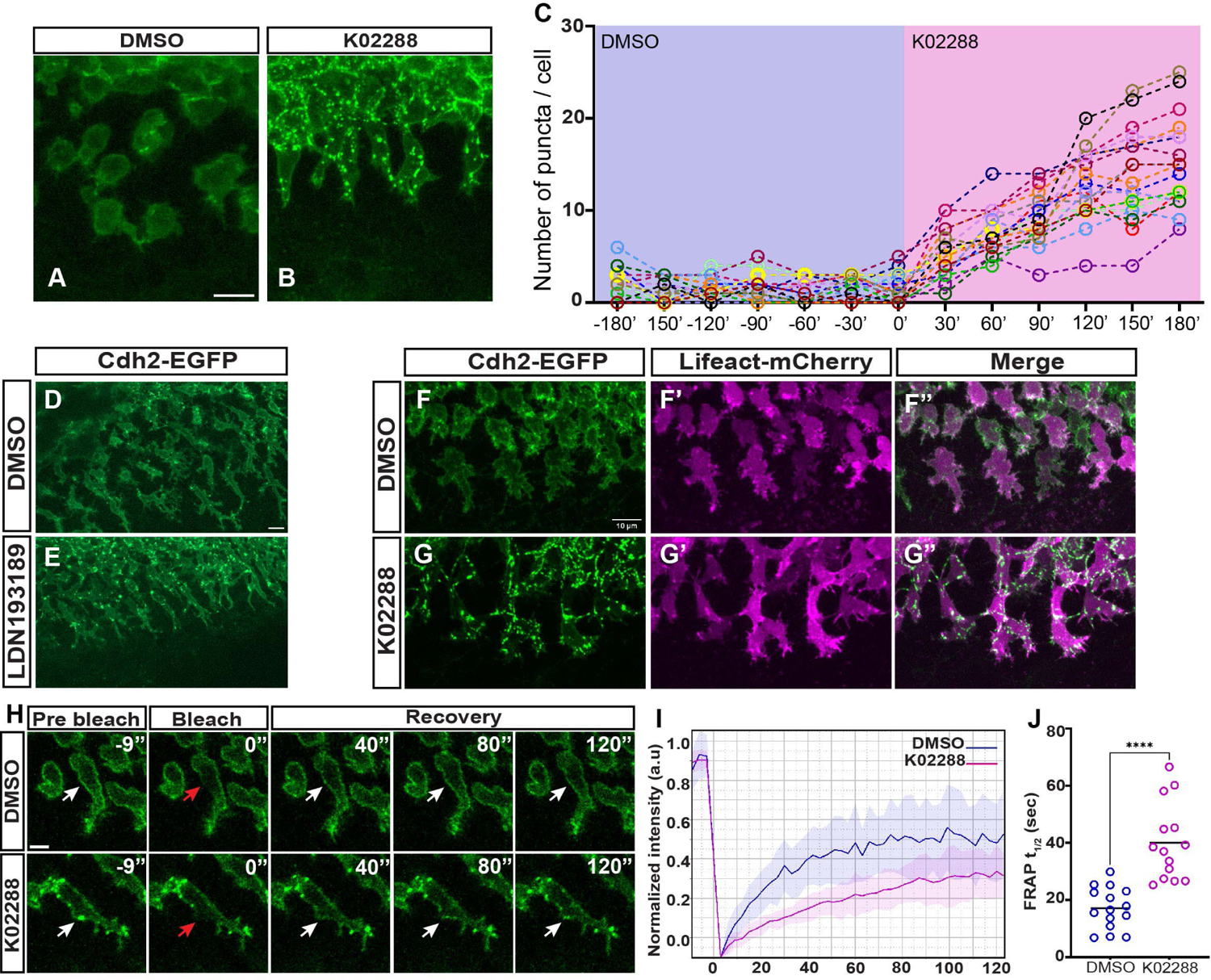
Inhibition of BMP signalling affects N-cadherin dynamics of fin mesenchyme cells. Fluorescence micrograph of TgBAC(*cdh2:cdh2-EGFP*) fin mesenchyme cells showing redistribution of Cdh2-EGFP under inhibition of BMP signalling (A,B). Counting the number of Cdh2-EGFP puncta in each cell reveal a progressive increase of Cdh2 aggregation over time upon initiation of BMP inhibition (C). A similar observation can be made upon treatment with LDN193189, an alternate pharmacological inhibitor of BMP signalling (D, E). Despite visible changes in the distribution of Cdh2-EGFP (F, G), no marked changes were seen in the mesenchyme cell actin cytoskeleton after treatment with K02288 (F’, G’) or in the association between Cdh2 and actin (F”, G”). During FRAP experiments on TgBAC(*cdh2:cdh2-EGFP*) embryos, we observed that Cdh2-EGFP puncta in K02288 treated embryos had a delayed fluorescence recovery compared to Cdh2-EGFP in control embryos (H). Plotting the fluorescence recovery curves (I) and fitting them with a single exponential recovery model showed a significant increase in the half-time of recovery (t_1/2_) in the K02288-treated embryos (J).

**Fig 6.**
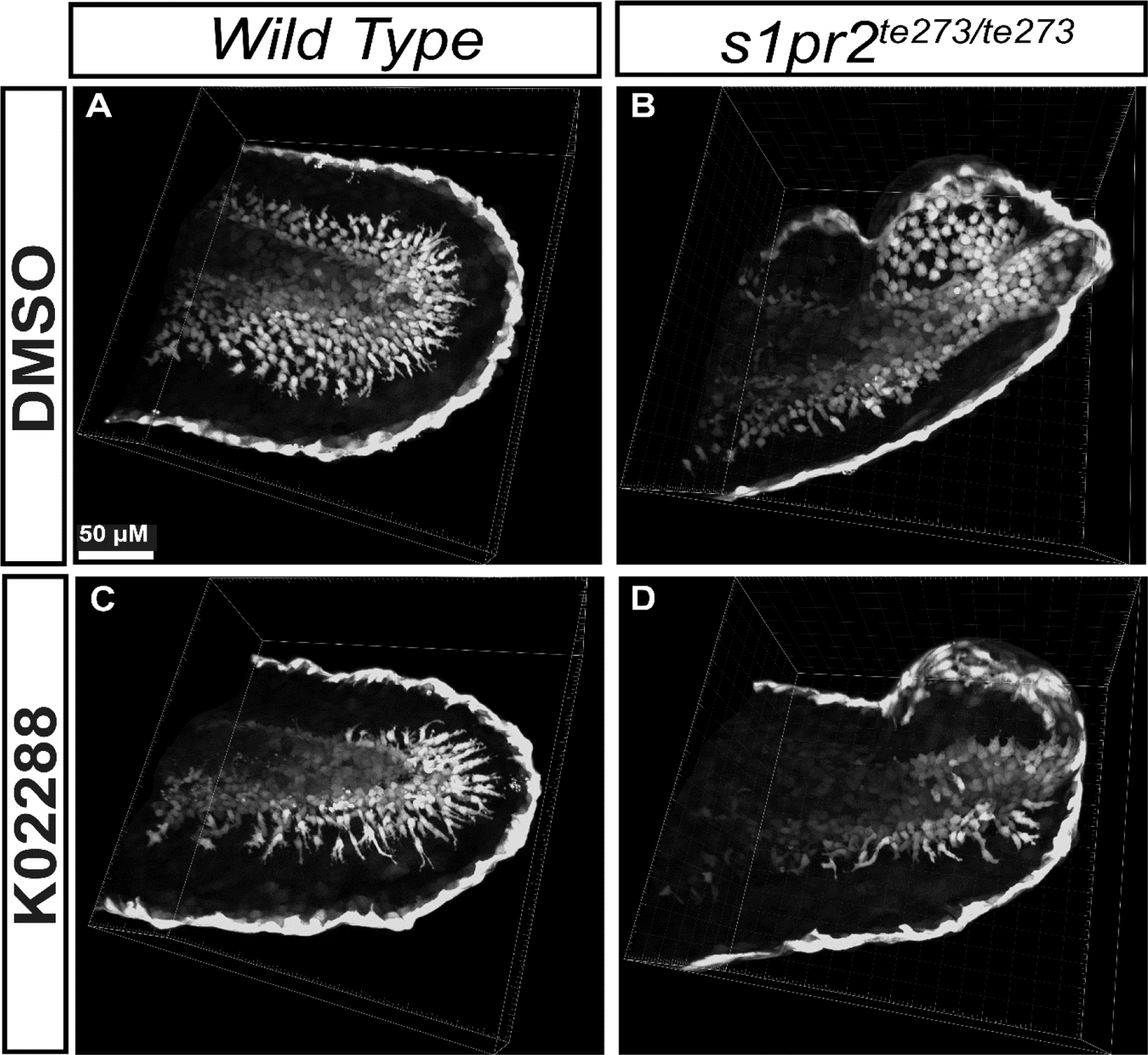
Three dimensional reconstruction of confocal micrographs of larval fin folds from either Wild-type or *s1pr2^-/-^* embryos treated with DMSO or K02288. Compared to WT fin folds (A), *s1pr2* fin folds under control conditions have ectopic fin blisters populated by non-polarized fin mesenchymal cells with normal degree of cell-cell contact (B). Upon K02288 treatment, the migration of fin mesenchymal cells in WT embryos is visibly reduced with increased cell-cell adhesion (C). K02288 treatment in *s1pr2^-/-^* embryos result in an additive effect wherein fin mesenchymal cells appear both non-polarized, exhibit inhibited migration and fail to delaminate from the mesoderm where they are specified, due to increased cell-cell attachment (D).

We also investigated cytoskeletal changes as a contributory factor in fin mesenchyme migration. To visualize cytoskeletal actin, we utilized the Tg(*tbx16l:LifeAct-EGFP*) transgene to drive LifeAct-EGFP expression in fin mesenchymal cells under the *tbx16l* promoter. However, we did not observe any significant changes in F-actin distribution under BMP-inhibited conditions. Previous work has linked the clustering of N-cadherin on cell membranes to an increased association with the actin cytoskeleton [57, 58]. As such, we looked at possible changes in N-cadherin-actin interaction usingTgBAC(*cdh2:cdh2-EGFP,crybb1:ECFP*);Tg(*tbx16l:LifeAct-mCherry*) double transgenic embryos. Actin fibers could be seen associated with both uniformly distributed Cdh2-EGFP in control cells as well as Cdh2-EGFP foci in K02288 treated embryos, but no notable difference could be observed in the actin association between the two groups (Figure 5F-F”, G-G”, Supplemental Movie 7).

Similar N-cadherin punctae have been observed in ventricular cardiomyocytes during cardiac trabeculation, and have been reported to display altered membrane dynamics [57]. To test for a similar possibility, we performed Fluorescence Recovery After Photobleaching (FRAP) experiments on the mesenchymal cell membranes of TgBAC(*cdh2:cdh2-EGFP,crybb1:ECFP*)embryos under treatment with K02288 or vehicle control. Evenly distributed Cdh2-EGFP molecules in control mesenchymal cells showed rapid fluorescence recovery after photobleaching (Figure 5H, Supplemental Movie 8). However, Cdh2-EGFP puncta in K02288 treated embryos failed to reform after photobleaching, with some degree of fluorescence recovery observed due to the surrounding Cdh2-EGFP molecules diffusing into the photo-bleached region (Figure 5H, Supplemental Movie 8). Quantitative analysis of fluorescence recovery curves also showed that Cdh2-EGFP under control conditions recovered significantly faster (half-time of recovery, t_1/2_ = 17.13 ± 7.05 sec) compared to BMP inhibited samples (t_1/2_ = 40.02 ± 12.98 sec) (Figure 5I, J), thereby indicating a role of BMP in regulating the membrane mobility of N-cadherin in fin mesenchymal cells.

Together, our data suggests that BMP signalling in the developing fin fold might regulate the cellular dynamics of N-cadherin in the fin mesenchymal cells during their migration, resulting in the organization of N-cadherin into membrane clusters upon BMP inhibition. However, unlike some previous reports, this redistribution of N-cadherin in the absence of BMP signalling appears to be independent of its association with the cytoskeletal network. This novel influence of BMPs on N-cadherin in mesenchymal cells may provide a mechanistic explanation for the observed reduction of migration and increased cell-cell attachment in the absence of BMP signalling.

## Discussion

Vertebrate limb bud morphogenesis is shaped by the interaction between AER-produced signals such as Wnt and FGF that coordinate to establish cell polarity in the limb mesenchyme and drive directional outgrowth [59, 60]. In zebrafish, a similar role is played by Slit-Robo signalling which regulates cell polarity and cell-ECM adhesion in the limb mesenchyme [61]. Studies in mice have also insinuated BMP signalling as a key player in proper limb development and digit identity determination, however, these studies did not seek to explore the cellular events or molecular mechanisms that are responsible for this physiological role of BMPs. Our work describes a novel function of BMP signalling in driving the morphogenesis of the zebrafish median fins, whose developmental mechanisms are evolutionarily linked to tetrapod paired appendages [62]. We show that BMP signalling is initiated in the median fin fold by AER-derived ligands activating their cognate receptors on cells of the fin mesenchyme, a transcriptomic profile consistent with previous observations in higher vertebrates [63, 64]. In addition to their shared expression domain in the AER, the three ligands each exhibited unique transcriptomic signatures in and around the fin mesoderm, suggesting distinctive supplementary functions during fin development. Similarly, even though both tested BMP receptors were broadly expressed throughout the fin mesoderm with a high degree of spatial overlap, their regions of most prominent expression appeared mutually exclusive, hinting at possible complimentary roles. The transcriptomic distribution of *id1* remained largely consistent throughout fin growth, with expression mainly observed in the fin mesoderm and its derivatives. Quantification of phosphorylated Smad 1/5/9 levels confirmed that this AER-to-mesoderm signalling relay sets up a gradient of BMP activity across the fin mesenchyme cells along the P/D axis, with the most distal cells closest to the ligand source in the AER displaying the highest degree of signalling activation.

In addition to the establishment of a BMP activity gradient, the dynamic scaling of such a gradient is also necessary for proper tissue morphogenesis. The development of the zebrafish pectoral fin is orchestrated by Smoc1, a BMP agonist that provides feedback regulation to scale the activity gradient and control fin outgrowth [41]. Similarly, the BMP activity gradient formed during Drosophila wing development is scaled by Pent, a secreted BMP regulator, whose loss is associated with reduced wing size and shorter gradient length-scale [65]. With the growth dynamic of the zebrafish median fin being comparable to that of the pectoral fin, it may be possible that similar feedback mechanisms are also at play in the median fin fold, thus providing exciting avenues for further research.

Disruption of BMP signalling in the developing fin, first using small molecule inhibitors of BMP receptors and then with a dominant negative BMP receptor, revealed its role in orchestrating the migration of the fin mesenchyme. In contrast to the vast majority of previously reported developmental roles of BMP in zebrafish, our findings are unique in that the pro-migratory role of BMP in the MFF appears to be non-morphogenetic, i.e. there is no effect on cell fate decisions. BMP has previously noted roles driving the chemotactic migration of mesoderm-derived cell lines *in vitro*, however, these observations have been attributed to the non-canonical Smad-independent signalling [17, 19]. Though our findings hinted at the non-involvement of the Smad-dependent canonical pathway, we also did not uncover substantial evidence of any causal non-canonical mechanisms. However, it is to be noted that our findings on non-canonical signalling are based on investigation of p-ERK and p-p38, which are only two of the many downstream effectors known to be associated with BMP signalling [13]. Thus, it is entirely plausible that other secondary pathways that were beyond the scope of our investigation could be responsible for directing mesenchyme migration. The elucidation of the exact nature of the signalling relay has proven challenging owing to the myriad of secondary effectors that can be recruited by an activated BMP receptor. One possible solution would be to conduct a high-throughput transcriptomic analysis of the migrating fin mesenchyme, but this prospect is compounded by the fact that the non-canonical pathway activates many downstream effectors in a transcription-independent manner via site-specific phosphorylation [13]. Thus, a combinatorial approach utilizing both phosphoproteomics and transcriptomics might be necessary to unravel the identity of the intracellular signalling cascade [66, 67].

Morphometric analysis of fin mesenchyme cells revealed significant alterations in cellular shape coincident with defects in migratory behavior. The morphological changes, particularly in cell spread area and circularity, are reminiscent of observations in cultured cells exposed to variable regimens of uniaxial cyclic stretch [68, 69]. On closer inspection of their migratory behavior, we found that mesenchymal cells also showed an increased tendency to remain in prolonged contact with their neighbors in the absence of BMP signalling. Taken together, these results allude to the possibility that mesenchymal cells subjected to BMP repression may undergo changes in cell-cell adhesion properties, which may then cause them to be subjected to increased tensile stress under the influence of migratory forces consequently leading to morphological deformations.

Our hypothesis regarding altered cellular adhesion gained further traction upon analysis of Cdh2-EGFP distribution in the fin mesenchymal cells of transgenic TgBAC(*cdh2:cdh2-EGFP*) embryos. Cdh2-EGFP could be observed in the membrane of migrating cells with generally higher levels observed in the proximal stationary cells compared to the distal migratory ones. The expression of Cdh2-EGFP across the fin mesenchyme cells of the P/D axis resembled a graded distribution that exhibited an inverse correlation with p-Smad1/5/8 levels in that same population. This served as the primary indication of a possible causal relationship between graded BMP signalling and Cdh2 levels in the fin mesenchyme. A similar negative regulation of N-cadherin by BMP signalling has been previously suggested during zebrafish gastrulation [54]. Further evidence surfaced when we compared the cellular distribution of Cdh2-EGFP in the absence of BMP signalling. Whereas Cdh2-EGFP was evenly distributed and highly mobile along the membrane boundary of migratory cells under control conditions, inhibition of BMP signalling led to clustering of Cdh2-EGFP molecules in bright puncta on the non-migratory cells which displayed reduced membrane mobility and fluorescence recovery upon irreversible photobleaching. Unlike some previous instances where the emergence of Cdh2 puncta were found to be associated with changes in the cytoskeletal network, we did not observe any changes in actin distribution during the perturbation of mesenchymal cell migration. Thus, it could be surmised that the clustering of Cdh2 on the membrane likely influences fin mesenchymal cell migration by affecting intercellular adhesion rather than the migratory machinery itself.

Previous research on vertebrate limb formation have identified the necessity of several signalling factors, yet certain key cellular events during this process remain poorly characterized. The outgrowth of the embryonic limb bud is one such crucial step during this developmental process and our findings propose a novel molecular mechanism in which graded BMP signalling in the zebrafish median fin fold promotes this event by coordinating fin mesenchyme migration via attenuation of intercellular adhesion. Though further research will be required to answer the questions that remain, this work provides a gateway into examining additional cellular processes that may be relevant during limb development as well as exploring previously overlooked non-morphogenetic roles of morphogenetic pathways.

## Materials and methods section

### Zebrafish lines and maintenance

Zebrafish were maintained at the fish facility at EMB (NTU) under standard conditions at 28°C on a 14-hour light and 10-hour dark cycle. Embryos were obtained through natural crosses, raised at 28°C in E3 media and staged according to Kimmel et al.[70]. Fish lines used were: wild-type AB/TL, sqet37Et (ET37) enhancer trap line [71], Tg(*BMPRE-AAV.Mlp:mKO2*)^mw40^ (hereafter referred to as BRE:mKO2) [46], Tg(*hsp70l:dnXla.Bmpr1a-GFP*)^w30^ [51] and TgBAC(*cdh2:cdh2-EGFP,crybb1:ECFP*) [56]. Tg(*tbx16l:nlsGFP-P2A-mCherryCAAX*), Tg(*hsp70l:dnbmpr1ba-EGFP,crybb:ECFP*) (hereafter referred to as dnbmpr1ba-EGFP),, Tg(*hsp70l:smurf1-P2A-mCherryCAAX, crybb:ECFP*) (hereafter referred to as hsp70:smurf1), Tg(*tbx16l:LifeAct-EGFP*) and Tg(*tbx16l:LifeAct-mCherry*) were generated during this study.

### Whole mount *in situ* hybridization

*In situ* hybridization was carried out on wild-type embryos fixed overnight in 4% PFA, according to previously described protocols [72]. The following sequences were used as anti-sense probes – *bmp2b* (NM_131360.2), *bmp4* (NM_131342.2), *bmp7b* (NM_001077146.2), *bmpr1ab* (NM_001004585.1), *bmpr1ba* (NM_131457.2) and *id1* (NM_131245.2). For certain samples in the ET37 background, immunostaining for GFP was performed after completion of the colour reaction. Antibodies used were mouse anti-GFP (B-2) (1:200; Santa Cruz; #sc-9996) and Alexa 488-conjugated donkey anti-mouse IgG (1:500, Invitrogen). Nomarski and fluorescent images were acquired on a Zeiss Axio Imager A2.

### Generation of DNA constructs for transgenesis

The DNA construct for heat-inducible overexpression of dominant-negative Type I BMP receptor (*pminiTol2-hsp70-dnbmpr1ba-EGFP-cryECFP*) was generated via Gibson assembly [73] using NEBuilder HiFi DNA Assembly Cloning Kit (NEB). In brief, a truncated dominant-negative form of the zebrafish *bmpr1ba* gene (NM_131457.2) (from aa1-aa233) was amplified from wild-type cDNA pool and assembled into a pminiTol2 backbone alongside a C-terminal EGFP tag and a *crybb:ECFP* transgenesis marker. The construct for overexpression of zebrafish *smurf1* (NM_001001943.1) gene (*pminiTol2-hsp70-smurf1-P2A-mCherryCAAX-cryECFP*) was generated in a similar fashion.

### Tol2 injections

pT3TS-Tol2 was linearized with SmaI (NEB) and used as template for *in vitro* transcription using mMESSAGE mMACHINE T3 Transcription Kit (Ambion) according to manufacturer’s instructions. Tol2 mRNA at a final concentration of 80 ng/µl was co-injected with DNA constructs into the cell of 1-cell stage wild-type zebrafish embryo. The concentration of injected DNA constructs was maintained between 20-25 ng/µl unless mentioned otherwise.

### Whole-mount IHC

All embryos for IHC were fixed overnight at 4°C in 4% PFA. After vigorous washing in PBS-Triton (PBS with 0.1% Triton X-100), embryos were permeabilized by washing for 1 hour in 0.5% PBS-Triton. Samples for phospho-p38 immunostaining were additionally permeabilized with 5µg/ml Proteinase K in PBS-Triton for 5 minutes before being washed in 0.5% PBS-Triton. Then samples were blocked for 2-3 hours at RT in primary blocking solution (10% NDS in 0.5% PBS-Triton) followed by overnight incubation at 4°C with primary antibodies diluted in primary blocking solution. After antibody removal, samples were thoroughly washed in PBS-Triton and blocked for 1-2 hours at RT in secondary blocking solution (5% NDS in PBS-Triton) followed by overnight incubation at 4°C with secondary antibodies diluted in secondary blocking solution. Following secondary antibody removal, samples were washed in PBS-Triton followed by equilibration in 80% glycerol and mounting on glass slides. Primary antibodies used were mouse anti-GFP (B-2) (1:200; Santa Cruz; #sc-9996), rabbit anti-phospho-Smad1/5/9 (1:200; Cell Signalling Technology; #13820), rabbit anti-phospho-p38 MAPK (T180/Y182) (1:150; Cell Signalling Technology; #9215), rabbit anti-phospho-p44/42 (Erk1/2) MAPK (T202/Y204) (1:200; Cell Signalling Technology; #4370). The follow secondary antibodies were sourced from Invitrogen: Alexa 488-conjugated donkey anti-mouse IgG (1:500), Alexa 647-conjugated donkey anti-rabbit IgG (1:400-1:500). Nuclei were counterstained with 1 µg/ml DAPI (Thermo Fisher Scientific).

### Expression induction by heat shock

Heat shocks for all embryos at 28 hpf were done at 40°C in a water bath for two 60-minute pulses with a 120-minute recovery period in between. For Tg*(1.5hsp70l:dnbmpr1ba-EGFP,crybb:ECFP)*, heat shock for gastrulating stages was done at 37°C in an air incubator for 60 minutes followed by screening of fluorescence 90 minutes into recovery. For Tg*(1.5hsp70l:smurf1-P2A-mCherryCAAX,crybb:ECFP)*, heat shock for gastrulating stages was done at 40°C in a water bath for 60 minutes followed by screening of fluorescence 120 minutes into recovery.

### Treatment with small molecule inhibitors

Small molecule antagonists of BMP signalling, K02288 and LDN193189 were purchased as lyophilized powder (Sigma-Aldrich) and resuspended in DMSO to a stock concentration of 20mM, then aliquoted and frozen at −20°C. For dose-response experiments, stocks were diluted in E3 media to indicated concentrations. For all remaining experiments including live imaging, K02288 was diluted in E3 media to a final working concentration of 40 µM.

### Fluorescence microscopy

Brighfield imaging was performed on a Zeiss Axio Zoom V16. Embryos were mounted in 1% low melting agarose containing 0.02% tricaine (buffered to pH 7.0) in petri dishes. Embryos were then overlaid with E3 media containing 0.02% tricaine (buffered to pH 7.0). Embryos for fluorescence imaging were mounted in a similar manner but imaging was performed on a Zeiss LSM800 upright confocal microscope. For time-lapse imaging, the agarose around the tail was removed to permit free movement during growth.

### Cell tracking

Drift-corrected time-lapse movies of cell migration were exported to Fiji [74] and selected cells were tracked over time using the ‘Manual Tracking’ plugin. Individual tracks were exported out as positional coordinates and custom codes in R were used to plot cell trajectories and wind rose plots.

### FRAP measurements

TgBAC(*cdh2:cdh2-EGFP,crybb1:ECFP*) embryos were mounted in 1% low melting agarose containing 0.02% tricaine in petri dishes and agarose around the tail was removed to permit free movement during growth. The embryos were staged and subsequently treated with 40 µM K02288 or vehicle control for 4 hours and then FRAP measurements were performed using a 40X water immersion objective on a Zeiss LSM800 upright confocal microscope. In brief, 3 pre-bleach images were acquired before a small circular region (∼4 µm in diameter) on the cell membrane was irreversibly photo-bleached by repeated scanning using the 488 nm laser line. Then, post-bleach images were acquired at 3 secs intervals for 120 secs. Data pre-processing including background subtraction, bleach correction and normalization as well as curve fitting was performed using easyFRAP [75] application. Half-time of recovery (t_1/2_) was calculated by fitting the recovery curves with a single exponential equation.

### Image processing

Image processing for immunostaining experiments was performed on Imaris 9.5 (Bitplane). Due to high background signals from the skin obtained during immunostaining of p-Erk1/2 and phospho-p38, surface segmentation and selective signal masking was performed to isolate the signal in the region of interest. For the p-Erk1/2 IF experiments, the GFP signal was used to segment the fibroblasts using the ‘Surface’ function in Imaris and all p-Erk1/2 intensity outside the segmented surface was set to zero, thereby isolating the p-Erk1/2 signal inside the fibroblasts. For phospho-p38 IF, the GFP signal was similarly used to segment the fibroblasts and the DAPI signal outside the segmented fibroblasts was set to zero. Having isolated the fibroblast nuclei, this signal was used to build a second surface and all phospho-p38 signal intensity outside the second surface was masked, thereby isolating the nuclear phospho-p38 signal in the fibroblasts. p-Erk1/2 and phospho-p38 quantification was performed on Imaris and exported.

### Statistical analyses

Difference between treatment groups were tested for statistical significance using Student’s unpaired two-tailed t-test with ‘p < 0.05’ considered significant. All statistical analysis was carried out on Prism 9 (Graphpad).

---

**Supplementary Figure 1.**
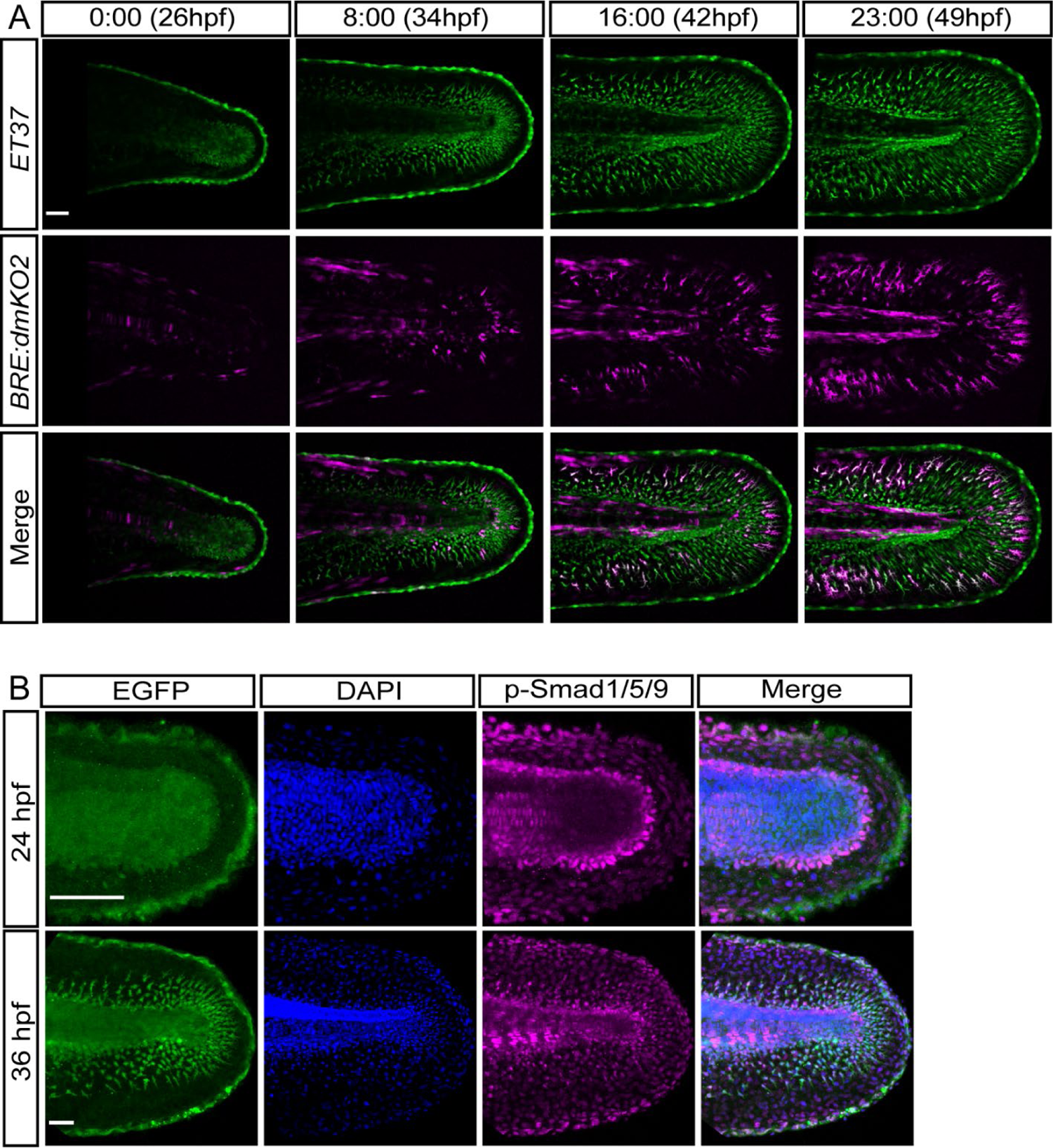
Extract from time-lapse movie showing migration of the fin mesenchyme and corresponding increase in BMP activity between 26 hpf and 49 hpf (A) phospho-Smad1/5/9 immunofluorescence shows BMP activity in the fin fold at 24 hpf and 36 hpf (B)

**Supplementary Figure 2.**
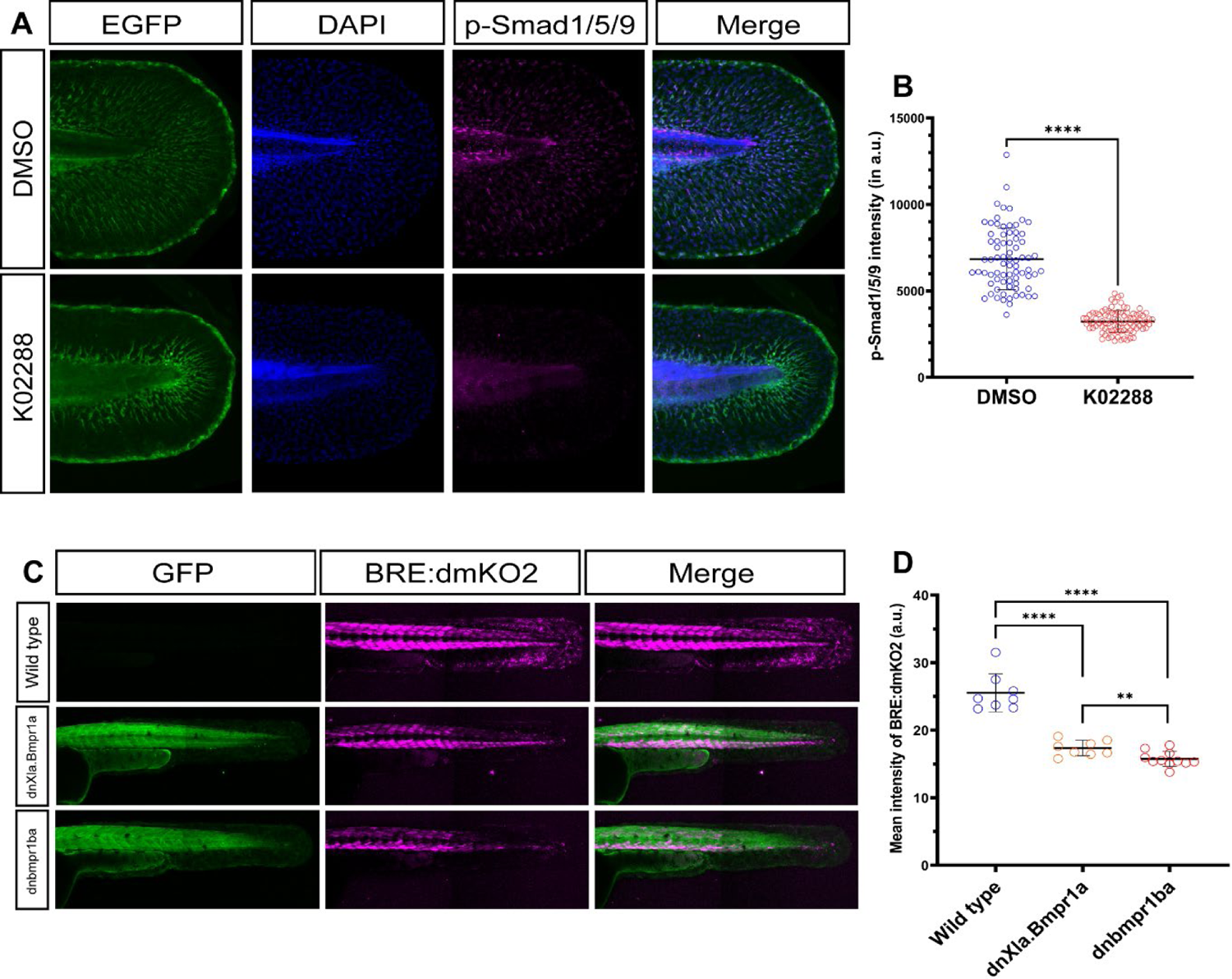
Immunofluorescence with pSmad1/5/9 antibody shows reduction of BMP activity in the fin mesenchyme upon treatment with K02288 (A, B). BMP reporter activity in the fin fold is significantly diminished upon over-expression of either dominant-negative BMP receptor (C, D).

**Supplementary Figure 3.**
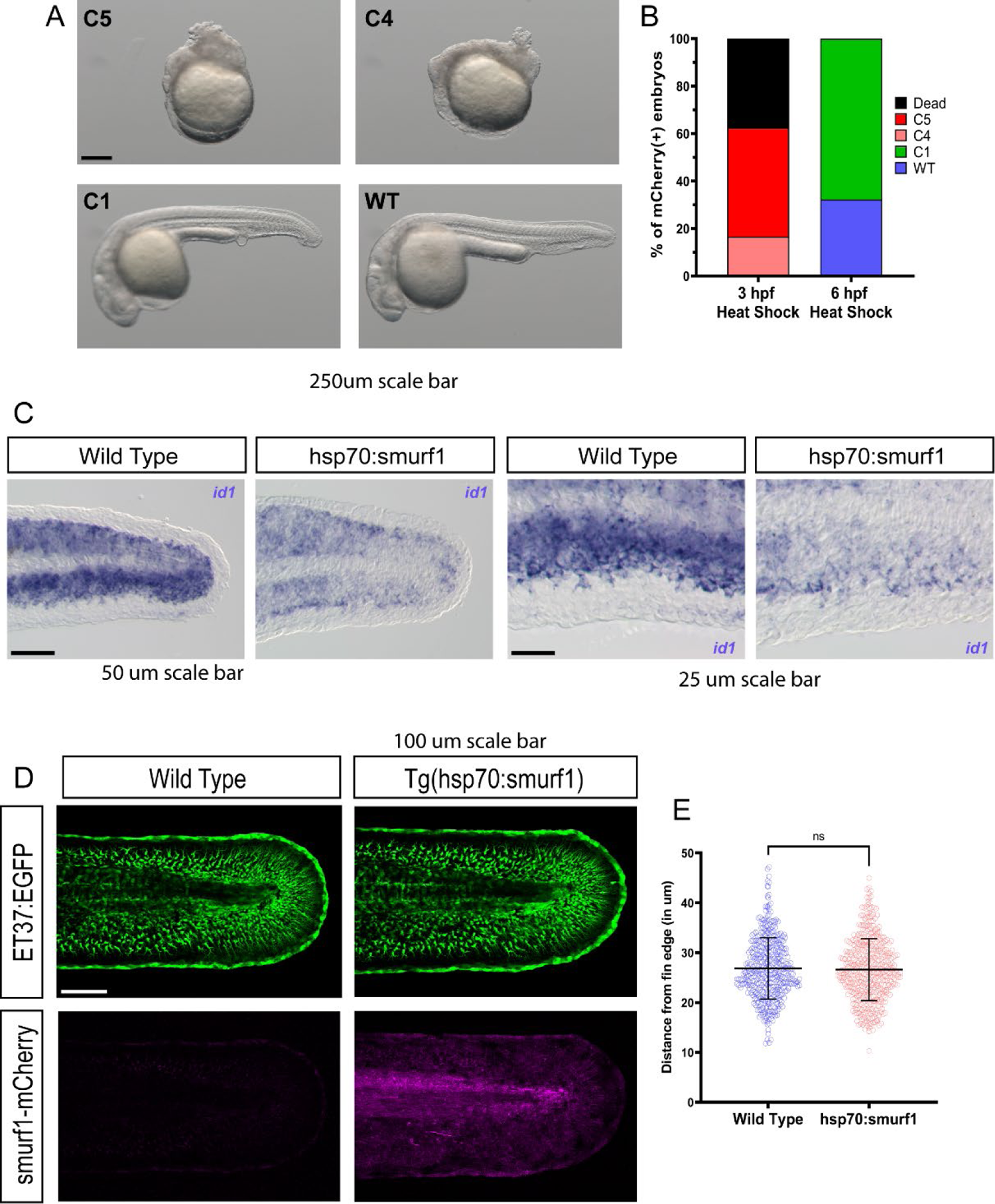
Transgene activation of hsp70:smurf1 at either 3 hpf or 6 hpf induced dorsalization of embryos when observed at 24 hpf (A). The severity of the phenotype depended upon the stage of transgene activation similar to observation made with dominant negative BMP receptors (B). Transgene expression at 28 hpf reduced *id1* transcript levels in the fin mesenchyme (C). However, Smurf1 over-expression in the fin fold did not have any significant impact on mesenchyme migration distance (D, E).

**Supplementary Figure 4.**
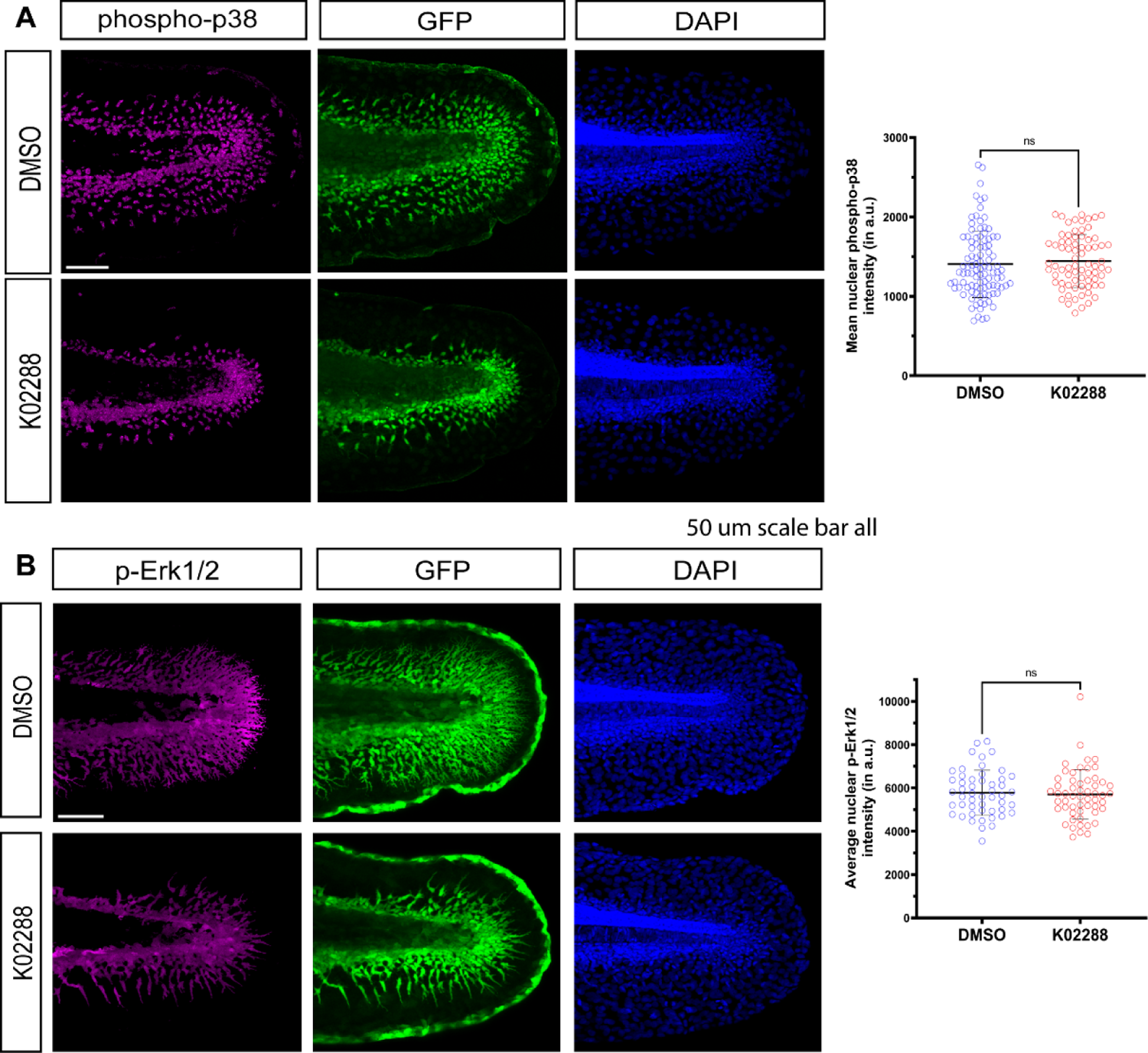
Upon treatment with K02288, mesenchyme migration was affected yet no significant changes were observed in p38 activity as measured by nuclear levels of phosphorylated p38 in the fin mesenchyme cells (A). Similarly, no changes in ERK activity could be detected when fin mesenchyme cells were stained with phospho-Erk1/2 antibody (B).

**Supplementary Figure 5.**
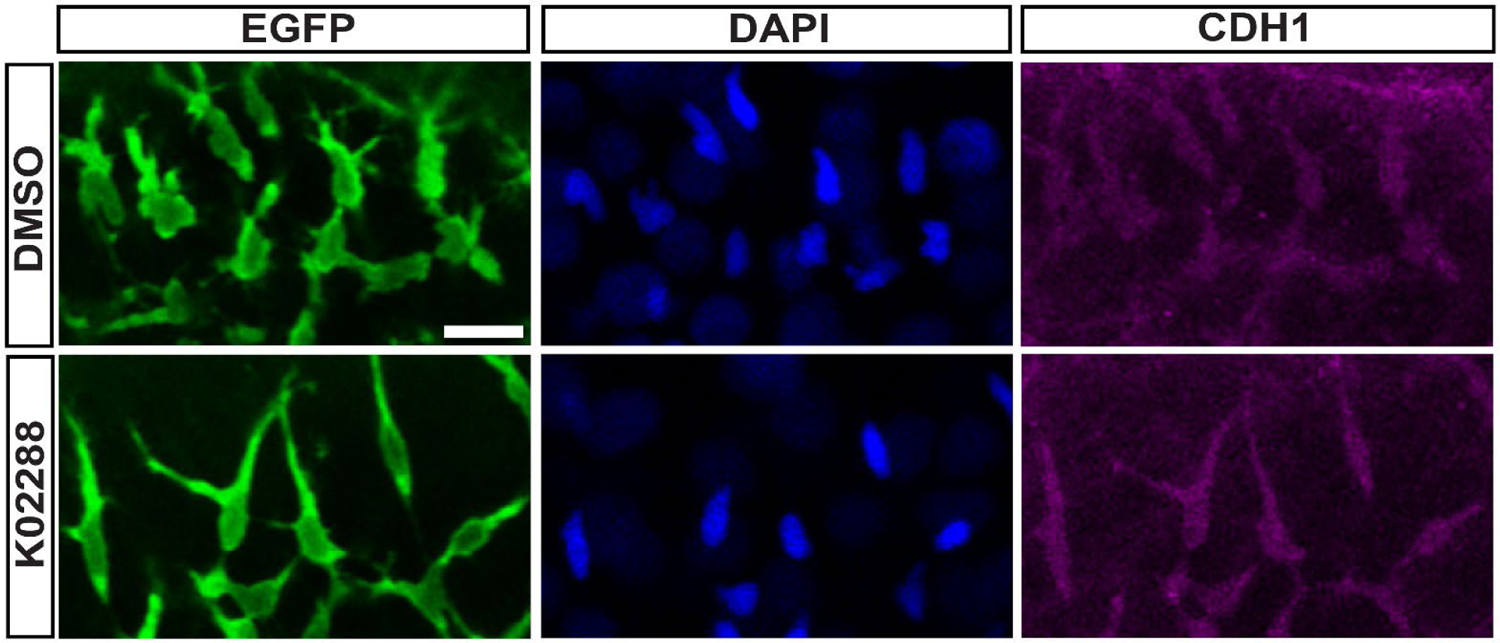
Immunofluorescence with anti-Cdh1 antibody did not show any discernable changes in E-cadherin expression in fin mesenchymal cells upon treatment with K02288.

**Supplemental Movie 1 - BMP activity dynamics in migrating fin mesenchyme cells.** Confocal time-lapse movie of the caudal fin of an Tg(*BRE:dmKO2)*;ET37 double transgenic embryo, imaged at 20X magnification (0.5x zoom) from 28 hpf for ∼21 hours at 10-minute intervals. The ET37-eGFP signal is shown in green, with the Tg(*BRE:dmKO2)* reporter in magenta.

**Supplemental Movie 2 - Migration of fin mesenchyme under control and BMP-inhibited conditions.** Epifluorescence time-lapse movie of the caudal fin of ET37 transgenic embryos under DMSO-treatment (left panel) or K02288-treatment (right panel). The embryos were treated with either DMSO or K02288 from 28 hpf and imaged at 10X magnification for ∼18 hour at 10-minute intervals.

**Supplemental Movie 3 - Intercellular interactions of fin mesenchyme cells under control and BMP-inhibited conditions.** High-resolution confocal time-lapse movie of the caudal fin of ET37 transgenic embryos under DMSO-treatment (top panel) or K02288-treatment (bottom panel). The embryos were treated with either DMSO or K02288 from 29 hpf and imaged for 90 minutes at 2-minute intervals under a 40X objective..

**Supplemental Movie 4 - Restoration of BMP signaling reverses the increased intercellular contact associated with reduced migration.** Confocal time-lapse movie of the caudal fin region of an ET37 transgenic embryo. The embryo was treated with K02288 from 28 hpf and imaged from 30 hpf onwards for 2 hours at 2-minute intervals, then the K02288 was washed out, replaced with DMSO and imaging continued for an additional 4 hours at 2-minute intervals. Arrows point to individual cells that detach from their neighbours and initiate migration upon removal of K02288.

**Supplemental Movie 5 - N-cadherin expression in migrating fin mesenchyme cells.** Confocal time-lapse movie of the tail region of a TgBAC(*cdh2:cdh2-EGFP,crybb1:ECFP*) transgenic embryo, imaged at 20X magnification (0.5x zoom) from 28 hpf for ∼21 hours at 10-minute intervals.

**Supplemental Movie 6 - Change in the cellular distribution of N-cadherin upon BMP inhibition.** Confocal time-lapse movie of the caudal fin in a TgBAC(*cdh2:cdh2-EGFP,crybb1:ECFP*) transgenic embryo, showing migrating cells in the ventral fin mesenchyme. The embryo was treated with DMSO from 29 hpf and imaged for 3 hours at 30-minute intervals under a 40X objective, then the DMSO was washed out, replaced with K02288 and imaging continued for an additional 3 hours at 30-minute intervals.

**Supplemental Movie 7 - Association of N-cadherin and F-actin in fin mesenchyme cells under control and BMP-inhibited conditions.** Confocal z-scan of the TgBAC(*cdh2:cdh2-EGFP,crybb1:ECFP*);Tg(*tbx16l:LifeAct-mCherry*) double transgenic embryos at 40x magnification under DMSO-treatment (top panels) or K02288-treatment (bottom panels). Images were taken at 32 hpf after 4 hours of treatment with either DMSO or K02288. The TgBAC(*cdh2:cdh2-EGFP,crybb1:ECFP*) reporter is shown in green with the Tg(*tbx16l:LifeAct-mCherry*) signal in magenta.

**Supplemental Movie 8 - Fluorescence recovery after photobleaching (FRAP) of Cdh2-EGFP under control and BMP-inhibited conditions.** Confocal time-lapse movie of the fluorescence recovery of photo-bleached fin mesenchymal cells in TgBAC(*cdh2:cdh2-EGFP,crybb1:ECFP*) transgenic embryos under DMSO-treatment (left panel) or K02288-treatment (right panel). The images were taken at 63x magnification at 32 hpf after 4 hours of treatment with either DMSO or K02288, at 3-second intervals with the first 3 frames representing pre-bleach fluorescence and last 40 frames being post-bleach recovery.

